# In vivo Imaging using Surface Enhanced Spatially Offset Raman Spectroscopy (SESORS): Balancing Sampling Frequency to Improve Overall Image Acquisition

**DOI:** 10.1101/2023.09.17.558110

**Authors:** Fay Nicolson, Bohdan Andreiuk, Eunah Lee, Bridget O’Donnell, Andrew Whitley, Nicole Riepl, Deborah Burkhart, Amy Cameron, Andrea Protti, Scott Rudder, Jiang Yang, Samuel Mabbott, Kevin M. Haigis

## Abstract

**Background and Rationale:** In the field of optical imaging, the ability to image tumors at depth with high selectivity and specificity remains a challenge. Surface enhanced resonance Raman scattering (SERRS) nanoparticles (NPs) can be employed as image contrast agents to specifically target cells *in vivo*, however, this technique typically requires time-intensive point-by-point acquisition of Raman spectra, thus hindering the real-time image acquisition desired for clinical applications. Moreover, traditional approaches involving Raman spectroscopy are limited in their inability to probe through tissue depths of more than a few millimeters. Here, we combine the use of “spatially offset Raman spectroscopy” (SORS) with that of SERRS in a technique known as “surface enhanced spatially offset resonance Raman spectroscopy” (SESORRS) to image deep-seated tumors *in vivo*. Additionally, by accounting for the laser spot size, we report an experimental SESORRS approach for detecting both the bulk tumor, subsequent delineation of tumor margins at high speed, and the identification of a deeper secondary region of interest with fewer measurements than are typically applied.

**Methods:** To enhance light collection efficiency, four modifications were made to a previously described custom-built SORS system. Specifically, the following parameters were increased: (i) the numerical aperture (NA) of the lens, from 0.2 to 0.34; (ii) the working distance of the probe, from 9 mm to 40 mm; (iii) the NA of the fiber, from 0.2 to 0.34; and (iv) the fiber diameter, from 100 µm to 400 µm. To calculate the sampling frequency, which refers to the number of data point spectra obtained for each image, we considered the laser spot size of the elliptical beam (6 × 4 mm). Using SERRS contrast agents, we performed *in vivo* SESORRS imaging on a GL261-Luc mouse model of glioblastoma at four distinct sampling frequencies: par-sampling frequency (12 data points collected), and over-frequency sampling by factors of 2 (35 data points collected), 5 (176 data points collected), and 10 (651 data points collected).

**Results:** In comparison to the previously reported SORS system, the modified SORS instrument showed a 300% improvement in signal-to-noise ratios (SNR). Glioblastomas were imaged *in vivo* using SESORRS in mice (n = 3) and tumors were confirmed using MRI and histopathology. The results demonstrate the ability to acquire distinct Raman spectra from deep-seated glioblastomas in mice through the skull using a low power density (6.5 mW/mm^2^) and 30-times shorter integration times than a previous report (0.5 s versus 15 s). The ability to map the whole head of the mouse and determine a specific region of interest using as few as 12 spectra (6 second total acquisition time) is achieved. Subsequent use of a higher sampling frequency demonstrates it is possible to delineate the tumor margins in the region of interest with greater certainty. In addition, SESORRS images indicate the emergence of a secondary tumor region deeper within the brain in agreement with MRI and H&E staining.

**Conclusion:** In comparison to traditional Raman imaging approaches, this approach enables improvements in the rapid detection of deep-seated tumors *in vivo* through depths of several millimeters due to improvements in SNR, spectral resolution, and depth acquisition. This approach offers an opportunity to navigate larger areas of tissues in shorter time frames than previously reported, identify regions of interest, and then image such area with greater resolution using a higher sampling frequency. Moreover, using a SESORRS approach, we demonstrate that it is possible to detect secondary, deeper-seated lesions through the intact skull.

## INTRODUCTION

In the field of medical imaging, the ability to detect deep-seated tumors in real time is both necessary and challenging. Utilizing light as an excitation source, optical imaging encompasses techniques including fluorescence, bioluminescence, optoacoustics, and Raman spectroscopy [1,2]. Optical imaging modalities can provide real-time, high-resolution visualization and characterization of structures and biological processes at the cellular and molecular level. This, together with its relatively low cost, high throughput, high spatial resolution, and ability to provide molecularly specific information, makes optical imaging modalities highly suitable for applications involving real-time imaging of cancer *in vivo* [3]. Currently, fluorescence based approaches lead the way in optical molecular- and intraoperative-imaging [2,4–7], however, fluorescence suffers from photobleaching and intrinsic autofluorescence [8]. Furthermore, fluorescent agents often exhibit broad and overlapping emission bands, which can restrict their utility in multiplexing applications [6].

Raman spectroscopy is a high-resolution, non-destructive optical molecular imaging strategy which relies upon the collection of inelastic light scattering following excitation with a laser source [9]. Its applicability in its intrinsic form (i.e. without the use of any contrast agent) has been demonstrated extensively in a number of biomedical applications including *in vivo* imaging [9–12] and image guided surgery in patents with grade 2 to 4 gliomas [13]. However, while such an approach successfully enables the discrimination between the different tissue types, (e.g. healthy or cancerous), intrinsic Raman spectroscopy is often associated with poor signal to noise ratios (SNR), long acquisition times, and typically relies on post-processing methods to deconvolute the spectra information, therefore hindering its potential for use for *in vivo* applications [14,15]. Nanoparticle (NP) based contrast agents (CAs) in the form of surface enhanced Raman scattering (SERS) NPs and surface enhanced resonance Raman scattering (SERRS) NPs have thus been employed to in part, circumvent the aforementioned limitations.

SERRS CAs are often composed of a core-shell structure which consist of a gold nanoparticle core functionalized with a resonant Raman reporter, which is then encapsulated in a silica shell [16–18]. By tracking their unique “fingerprint” spectra, SERRS CAs can be used to image tumors by targeting cells with high specificity *in vivo* (e.g. using antibodies [19–22] or peptides [23,24]). Non-targeted SERRS CAs have also been shown to accumulate in solid tumors due to the enhanced permeability and retention (EPR) effect [15,17,25–28] with several reports demonstrating the reliable detection of a large variety of different tumor types *in vivo using* SERRS CAs, including microscopic (< 100 µm) and their microscopic extensions using SERRS NPs [16,17,19,20,23,29,30]. Unlike fluorescence imaging, SERRS imaging allows the visualization of a large number of molecular markers within the same tumor sample simultaneously due to the unique “fingerprint” spectra of each “flavor” of SERRS CAs [6,27,31,32].

Despite several reports in the literature demonstrating the applicability of SERRS CAs to cancer imaging, the approach is largely limited to surface-based lesions due to the principal of light scattering in turbid media, including biological tissue. Spatially offset Raman spectroscopy (SORS) is an emerging optical molecular imaging approach which takes advantage of light-scattering properties of tissue to enable the visualization of deep-seated region(s) of interest (ROI) such as malignant tumors [33], while also suppressing autofluorescence contributions from the tissue [34,35]. The approach capitalizes on the statistical phenomenon whereby photons originating from deeper within a medium exhibit a greater propensity for lateral migration away from the point of illumination compared to their surface-level counterparts [34,35], thus light is scattered in multiple directions before it is gathered by the collection optics [35–37]. Surface enhanced spatially offset resonance Raman spectroscopy (SESORRS) describes the use of SORS to detect the accumulation of SERRS CAs in an ROI and offers the potential to image cancer with high contrast *in vivo* at depths far superior to what can currently be achieved using other optical imaging approaches [35,36,38,39]. SESORRS has been shown to successfully detect the accumulation of SERRS CAs through depths of up to 14 cm [40–42]. Recently, SESORRS has been applied to a number of biomedical applications including the detection of neurotransmitters [43,44], *ex vivo* breast cancer tumor models and phantoms [40,45,46], and the detection of glioblastoma *in vivo* [26].

The scientific literature contains studies investigating the application of SERS (Surface-Enhanced Raman Scattering) and SERRS (Surface-Enhanced Resonance Raman Scattering) nanotags for intra-operative cancer imaging *in vivo*, employing conventional confocal Raman spectroscopy techniques. Additionally, SESORS (Surface-Enhanced Spatially Offset Raman Spectroscopy) and SESORRS has been explored for probing clinically relevant depths, both *ex vivo* and in phantom models. [40,41,45,46]. However, there are very few reports on the use of SESORRS for imaging of disease *in vivo* [33] and none on the application of sampling frequency approaches to increase the overall speed of image acquisition over a ROI in a biological subject. Here, we report for the first time, an experimental SESORRS approach for detecting both the bulk tumor and subsequent high-speed delineation of tumor margins, with fewer measurements than typically applied. This was achieved through optimization of a custom-built SORS probe which generated significantly SNRs in comparison to the previous set up. The custom-built collection probe was used to image the accumulation and uptake of Raman active SERRS nanotags in a GL261 mouse model of glioblastoma (GBM). Previously, our group demonstrated the first application of the SESORRS approach for the imaging of GBM through the intact skull however this approach used long acquisition and accumulation times, higher laser powers, and did not consider sampling frequency [33]. By improving our collection efficiency through the engineering of a custom built SORS collection probe, significant reduction in integration time, and utilization of sampling frequency approaches, we report the successful detection of GBM though the intact skull in as little as 6 seconds. In doing so, we demonstrate a means to detect a ROI and then delineate the ROI with greater certainty and improved resolution. In addition, SESORRS imaging detects the emergence of a secondary ROI deeper within the brain, in agreement with MRI and H&E staining thus suggesting the suitably of SESORRS as intraoperative imaging approach for the detection and imaging of deep-seated tumors.

## MATERIALS AND METHODS

### Reagents

All chemicals were purchased from Sigma-Aldrich and used as received unless otherwise stated. Paraformaldehyde (16%) was purchased from Thermofisher and diluted to 4% in phosphate buffered saline (pH 7.1). Deionized water (DI), 18 MΩ⋅cm, was used in all experiments.

### SERRS nanotag synthesis

SERRS nanotags were synthesized according to a protocol based on recent publications by our group, with minor modifications as detailed below [15,33].

### Synthesis of gold seeds

5 nm gold seeds were synthesized in accordance with our previously described protocol [47]. 2 ml of 25 mM aqueous solution of HAuCl_4_ were added to 200 ml of DI water. Afterwards, 6 ml of freshly-prepared ice-cold 100 mM NaBH_4_ aqueous solution was added to the mixture under stirring and left overnight at room temperature before use.

### Synthesis of bare gold nanostars

The seed-mediated synthesis of gold nanostars was performed in a cold room (+4 °C). To 1800 ml of 200 mM solution of ascorbic acid, 5 ml of gold seeds from previous step were added, followed by 2.5 ml of 400 mM solution of HAuCl_4_. The obtained deep blue solution was rapidly transferred to 50 ml Falcon tubes and centrifuged at 0 °C and 3300 × g for 20 mins. After centrifugation, the transparent supernatant was removed and the liquid pellets on the bottom of the tubes were combined in a 15 ml Slide-A-Lyser G2 dialysis cassette with 20k MWCO and dialyzed in DI water. The water was changed daily over the 72-hour timeframe.

### Simultaneous Raman reporter attachment and silica encapsulation

To a 50 ml falcon tube (tube A), 1.2 ml of tetraethyl orthosilicate, 30 ml of isopropanol and 0.16 ml of 20 mM IR780p dye solution in DMF were added together. To a 15 ml falcon tube (tube B), 4 ml of 1 nM bare nanostars and 9 ml of ethanol were added. Immediately before combining tube A and tube B, 0.6 ml of ammonium hydroxide solution (28% aq) was added to tube B and tube B was shaken by rotation. The contents of tube B were added rapidly to tube A and the resulting mixture was left on a shaker for 20 min at room temperature. Next, tube A contents were divided equally between two 50 ml falcon tubes, which were each then filled with ethanol to 50 ml to quench the reaction, and then centrifuged at 3300 × g at 0 °C for 20 min. The dark green supernatant was removed, leaving ∼0.5 ml of solutions with dark blue liquid pellets on the bottoms of the tubes. The pellets were sonicated to fully homogenize the solution and transferred to two 1.5 ml Protein LoBind Eppendorf tubes, which were then filled with ethanol. The tubes were centrifuged at 10000 × g at room temperature for 5 minutes and the supernatants were discarded. The resulting pellets were then resuspended in 1 ml of ethanol and subjected to sonication. Ethanol washing was repeated three more times.

### Surface modification with PEG

To each 1 ml solution of nanostars resuspended in ethanol from the previous step, 0.1 ml of (3-mercaptopropyl)-trimethoxysilane and 0.04 ml of ammonium hydroxide solution (28% aq) were added and the reaction mixture was left on a shaker for 1h at room temperature. Then they were centrifuged at 10000 × g at room temperature for 5 minutes, washed with ethanol three times as described above, washed once with water, and then each resuspended in 1 ml of 10 mM HEPES buffer pH 7.3 to obtain thiolated nanostars solutions. PEG5k-Mal (Sigma-Aldrich 63187, CAS 99126-64-4) was dissolved in anhydrous DMSO at the concentration of 20mg/ml. 0.1 ml of this DMSO solution of PEG was added to each of the two solutions of thiolated nanostars in 10 mM HEPES buffer pH 7.3 to create a ∼200,000:1 PEG/NPs ratio. The Eppendorf tubes were left on a shaker for 1h at room temperature. The nanoparticles were spun down and washed 3 times with water and then resuspended in 10 mM HEPES buffer pH 7.3. Prior to injection, the nanoparticles were concentrated to 2 nM by spinning down the pellets, combining them, and resuspending in 1 ml of HEPES.

### SERRS nanoparticle characterization

Transmission electron microscopy (TEM) was performed using a JEOL 1200EX Transmission Electron Microscope at 80 kV. The sample was prepared as follows: 1 µL of ∼0.1 nM nanoparticles solution was left to dry onto a TEM grid (CF300-Cu, Electron Microscopy Sciences) for 30 min. Zeta potential measurements were performed using Zetasizer Nano ZS (Malvern) on particles dispersed in 10 mM HEPES buffer (pH 7.3). Nanoparticle tracking analysis (NTA) NS500 (Malvern) was used to determine nanoparticle concentration. UV2600 (Shimadzu) was used for optical characterization of nanoparticles.

### Animal models

All animal experiments were approved by the Institutional Animal Care and Use Committees of Dana-Farber Cancer Center (protocol #08023). GL261-Luc cells were used to generate the tumor bearing mice. Briefly, 8 week-old BALB/cJ mice were injected intracranially with 100,000 GL261-Luc cells into the left striatum. Tumor incidence and size was determined by MRI (Bruker BioSpec 7T/30 cm USR horizontal bore Superconducting Magnet System, Bruker Corp., Billerica, MA) 2 weeks after injection of GL261-Luc cells.

### Custom-built SORS collection probe

All SORS *in vivo* measurements were carried out using a custom built SORS collection probe.

### Custom-built SORS imaging system

All SORS measurements were carried out using a system built in house, the design of which was based on previous reports in the literature [45, 59, 60]. A 785 nm laser (Innovative Photonics Solutions) was coupled to a fiber optic probe (Innovative Photonics Solutions) and the lens was removed from the probe to enable the delivery of a diffuse collimated beam to the sample surface. Both the excitation and collection probe were mounted on individual *xyz* translational stages (Thorlabs) and a rotation mount was used to deliver the excitation light to the sample at a 45° angle. The incident beam was set to intersect the sample surface at a 45° angle and was therefore elliptical, with the shorter diameter being 4 mm and the longer 6 mm. The scattered Raman light was either collected by a previously reported, commercially available collection probe [33] or a custom built collection probe which was built to increase light collection efficiency (**Figure 1a**). The custom probe was designed as follows: The generated scattered Raman light was collected using a 0.34 NA lens (Thorlabs) and passed through two 785 nm dichroic beam-splitters (Semrock) angled at 45° mounted in cage cubes (Thorlabs). Light was then passed through a lens tube, an iris, and then a 785 nm longpass filter (Semrock) and an additional iris. The light was then passed through an additional 785 nm longpass filter (Semrock) positioned at 2°, a collimator, and then a 400 μm, 0.34 numerical aperture (NA) core fiber (Thorlabs) where it was delivered to a high throughput f/2 spectrometer (Innovative Photonics Solutions), to collect the scattered Raman photons. The collection probe was mounted perpendicularly to the sample surface and had a working distance of 40 mm. The spatial offset was controlled by translating laser beam away from the focal point of the collection probe in the region of a few mm (Δx). This directed the beam at an appropriate point on the sample, e.g., plastic or mouse. All samples were positioned on a third translational *xy* stage (Thorlabs) to allow freedom to move the sample without impacting the alignment between the excitation and collection probes. The SORS system using the custom-built collection probe is shown in **Figure 1b**.

**Figure 1:**
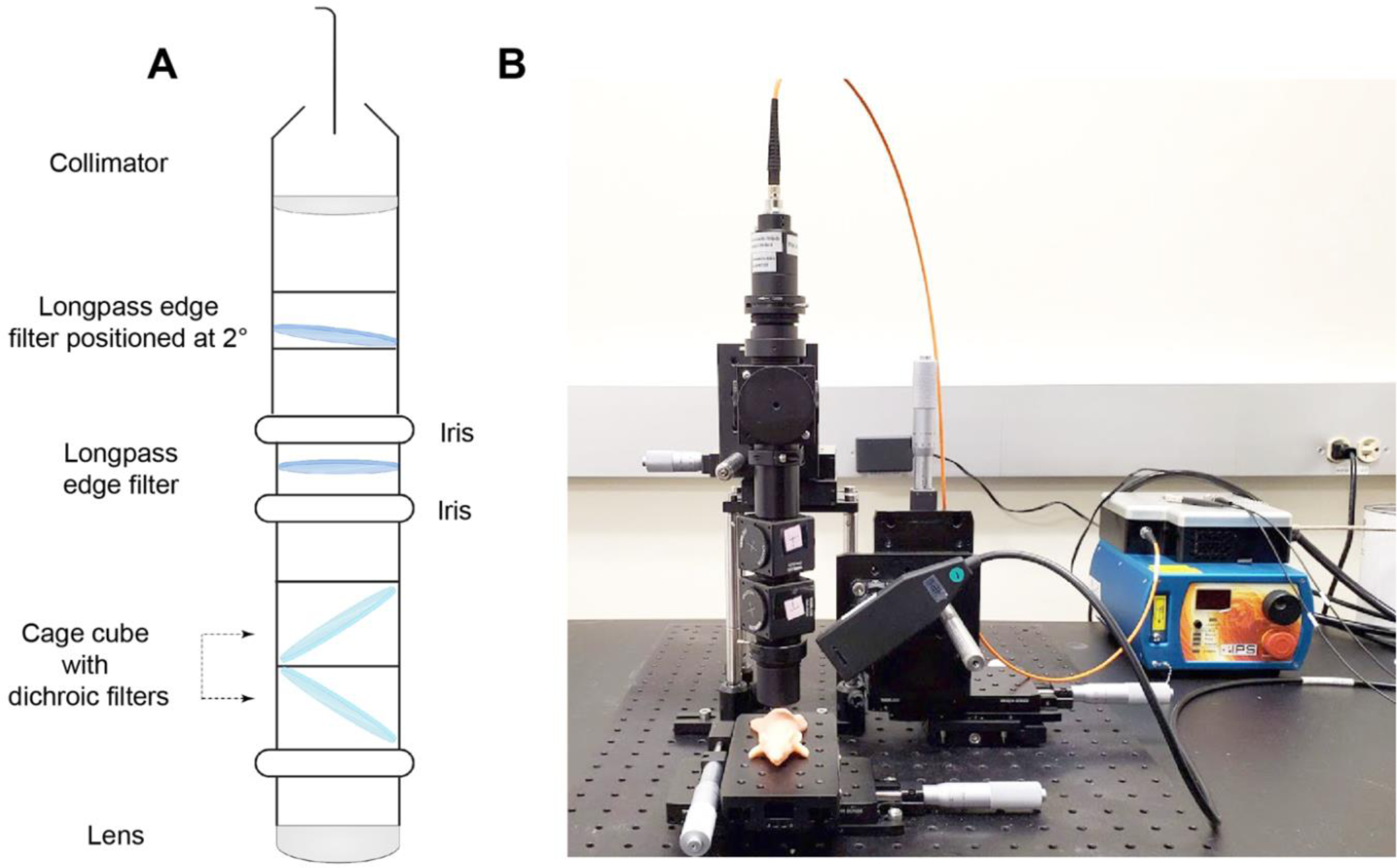
SORS Set-up using the custom-built probe. (A) Diagram describing the internal optics of the custom-built SORS collection probe. (B) In-house SORS set-up using the custom-built probe with an optical mouse phantom placed on the *xy* stage. Incident light is delivered at 45° to the sample surface and light is collected through the custom-built collection probe.

### SORS imaging parameters

Acquisition times and laser power were varied depending on the sample under study and are outlined below. Laser output was measured with a handheld laser power meter (Thorlabs). For SORS measurements using plastic phantoms a laser power of 500 mW was used. For brain tumor phantoms and animal models a laser power of 6 mW/mm^2^ at the sample surface was applied.

### SORS optimization measurements using plastic calibration standards

The efficiency of the custom built SORS collection probe in comparison to the previously used commercially available probe was evaluated using plastic calibration standards [33]. Polytetrafluoroethylene (PTFE) and pink polypropylene (PP) sheets were purchased from a plastic retailer. PTFE (5 cm (length) × 2 cm (width) × 2 mm (thickness)) was placed on the stage and varying numbers of PP sheets (length 5 cm × width 2 cm × thickness 2 mm) were then placed on top of the PTFE to act as a barrier (**Figure S1A**). Their corresponding Raman spectra are shown in **Figure S1B and C**. To increase the thickness of the barrier, further PP sheets were added. Measurements were carried out using a 4 second integration time, one accumulation, 500 mW, 785 nm laser. The spatial offset was controlled by moving the *xyz* translation stage coupled to either of the collection probes (commercially available or custom built) in the *x* direction.

### SORS optimization measurements using brain tumor phantoms

A brain tumor phantom was prepared using the head of a healthy 12-week-old-euthanized mouse. The brain was removed, and a small piece of PTFE (5 × 5 mm) was placed under the skull and glued in place. The skull was then filled with 1% agarose gel to create a phantom [47]. Unlike previous reports, the skin was not removed from the skull in order to create a more realistic phantom. Paraffin wax was melted using a heat plate and poured into a small plastic weigh boat. After the wax had partially cooled, the mouse’s head was positioned atop the semi-solid paraffin, which served as a mold for the base of the rodent’s cranium. The weigh boat containing the molded paraffin wax was then fixed in position on the *xy* translational stage. This allowed for consistent measurements of the mouse’s head over multiple days, ensuring that the head was positioned identically to the previous day. SORS measurements were carried out with the custom-built system at spatial offsets of 1, 1.5, 2, 2.5, and 3 mm. Again, the spatial offset distance was controlled by moving the *xyz* translation stage in the *x* direction. Measurements were acquired using different sampling frequencies (par sampling, 2x oversampling, 5x oversampling, and 10x oversampling). Spectra were obtained using a 0.5 second integration time, one accumulation, 6 mW/mm^2^, 785 nm. The step size used for each Raman map was dependent on the sampling frequency applied. Step sizes of 6 mm × 4 mm, 3 mm × 2 mm, 1.2 mm × 0.8 mm and 0.6 mm × 0.4 mm were used for a par-sampling, 2x oversampling, 5x oversampling, and 10x oversampling respectively. Step sizes for each sampling frequency were determined by the laser spot size (6 mm × 4 mm) and described in **Table 1**.

**Table 1:** Summary of step sizes applied to each sampling frequency and the total number of data points collected across a 12 × 12 mm area.

| Sampling frequency | Step-size in x | Step-size in y | Total number of data points |
| --- | --- | --- | --- |
| Par | 6 | 4 | 12 |
| Oversampling x2 | 3 | 2 | 35 |
| Oversampling x5 | 1.2 | 0.8 | 176 |
| Oversampling x10 | 0.6 | 0.4 | 651 |

### Imaging of GBM using SESORS

Prior to imaging, mice were shaved using an electric shaver and residual hair removed using hair removal cream. Each mouse was placed onto the *xy* translational stage with a range of 50 mm in each direction and moved in varying step sizes specific to each sampling frequency to acquire pointwise spectra. Mice (n=3) were injected via the tail vein with 2 nM SERRS CAs suspended in HEPES. Imaging was performed approximately 18 hours post injection of SERRS CAs.

### Biodistribution studies of SERRS CAs

Biodistribution analysis was performed to determine the fate of the SERRS nanotags *in vivo*. GBM-bearing mice (n=3) were injected with 2 nM of SERRS contrast agents (100 μL). Following administration, mice were euthanized using CO_2_ asphyxiation approximately 18 hours post injection. For each mouse (n = 3), tissues were harvested, weighed, and homogenized. Tissue homogenates were placed in 384-well plates. Raman images of the plates were acquired using 10% laser power (785 nm), 1 s acquisition, 5× objective (LabRAM HR Evolution (HORIBA Scientific. Spectra were baseline corrected, normalized to the maximum peak intensity, and the peak intensity at 947 cm^−1^ was plotted as a combination surface/contour false color 2D heat map. The peak at 947 cm^−1^ was selected since it is the strongest peak in the SERRS spectrum.

### Histology

Intact GBM-bearing brains were fixed in 4% paraformaldehyde (PFA) for 24 hours, transferred to 70% ethanol, and then embedded in paraffin. Each brain was sliced in the same orientation as the MRI and 5 μm-thick sections were stained with hematoxylin and eosin (H&E). Images were acquired using an Olympus BX-UCB slide scanner with a 20× objective and processed using OMERO software.

### Data processing

All spectra were processed using MATLAB software (version 2022b, The MathWorks) and LabSpec (version 6.6.2.7, HORIBA Scientific). SNRs were calculated by baselining the spectra and dividing the peak height of PTFE at 739 cm^−1^ by the intensity of the noise across 1799 cm^−1^ to 1899 cm^−1^. Processing of individual reference spectra involved truncating, and baseline correcting the spectra, followed by Savitzky–Golay smoothing. For the creation of false color 2D heat maps using brain tumor phantoms, spectra were truncated, baseline corrected, and smoothed using Savitzky–Golay filtering using MATLAB. The intensity of the peak at 739 cm^−1^ was then plotted as a combination surface/contour false color 2D heat map. For *in vivo* SESORRS images, spectra were truncated, baseline corrected using a polynomial fit and normalized using standard normal variate method in LabSpec. Principal component analysis (PCA) was then applied using the MVA-EVRI toolbox in LabSpec. Principal components (PC) 2 and 3 were folded to produce the spectral image. The intensity of peak height over the region of 898 cm^−1^ to 990 cm^−1^ was plotted and the data were extracted and transferred to MATLAB where they were plotted as combination interpolated/bilinear false color 2D heat maps.

In addition, we developed an in-house Python-based GUI for processing heat maps following PCA. The tool automatically identifies pixels in an image where the RGB values are all below 100, indicating they are a shade of black, and replaces them with a transparent pixel. This is achieved without altering the original dimensions of the heatmap. The process involved loading an image, applying the transparency adjustment, and saving the processed image. The GUI provides a user-friendly interface to choose an image, make black pixels transparent, and save the processed image. The code leverages the tkinter library for GUI creation and the Python Imaging Library (PIL) for image processing.

## RESULTS AND DISCUSSION

Our previous report of the use of SESORRS for the *in vivo* imaging of disease involved a commercially available collection probe to collect the scattered spatially offset photons. Although we successfully demonstrated the suitability of the approach, there were several limitations. Limitations included a short working distance from the sample surface and poor SNR, which in turn meant we had to utilize longer acquisition times and higher laser powers that reduced the applicability of our approach to *in vivo* imaging applications. To address these limitations and increase light collection efficiency, we therefore built our own SORS collection probe, as described in the methods section. In comparison to the commercially available probe, we applied four optimizations to our custom-built probe: (i) numerical aperture (NA) of the lens (0.2 to 0.34), (ii) working distance of the probe (9mm to 40mm), (iii) NA of the fiber (0.2 to 0.34), and (iv) fiber diameter (100µm to 400µm). The internal optics of the collection probe are described in **Figure 1A** and the final probe is shown in **Figure 1B**. To evaluate the efficiency of the new custom-built probe in comparison to the previously used probe, we used plastic calibration standards. Sheets of pink PP were placed on top of a PTFE layer to create a barrier with the aim of detecting the PTFE analyte through the PP barrier (Figure S1). Spectra of PTFE through PP thicknesses of 8, 14 and 20 mm of PP were acquired using the commercially available and custom-built SORS systems with spatial offsets of 0 to 10 mm (1 mm increments) The peak intensity of 739 cm^−1^ obtained through these thicknesses using either probe is shown in (**Figure 2A-C**). The 739 cm^−1^ peak was chosen, since it represents the most intense peak in the PTFE spectrum (**Figure S1B**). In both probe configurations, as the thickness of the polypropylene (PP) barrier increases, a larger spatial offset is necessary to achieve the highest polytetrafluoroethylene (PTFE) signal through the increased thicknesses. Notably, the custom-built probe consistently produces a higher signal than the commercially available probe across all tested thicknesses and offsets. **Figure 2D-F** describes the SNRs of each probe at 0 – 10 mm offsets through the same three thicknesses of plastic (8, 14 and 20 mm). As expected, the custom probe generates significantly higher SNRs through each of the three thicknesses. **Figure 2C** shows similar contribution of signal using the commercially available probe through 20 mm of PP at 0 – 10 mm offsets which is attributed to noisy spectra (i.e., the PTFE cannot be detected). This is confirmed in **Figure 2F**, which shows that the SNR for the commercial probe is consistently below 3. In contrast, our custom-built probe demonstrates the ability to detect PTFE through 20 mm of PP. As the thickness of the PP barriers increases, there is a notable reduction in both the SNRs and the relative intensity of the PTFE peaks. This decrease can be attributed to photon migration and energy loss, regardless of whether a commercially available probe or a custom-built probe is used. Furthermore, the data clearly indicates that as the PP barrier thickness increases, larger spatial offsets are necessary in order to achieve a significant spectral contribution from the PTFE. By increasing the spatial offset – specifically, the distance between the excitation spot and the collection point – it becomes possible to clearly discern the spectral contribution of PTFE through the PP barrier. These results indicate that our custom-built collection probe, which replaces the commercially available probe, has effectively improved our SORS system. We have achieved these enhancements due to increased Raman throughput and higher SNRs.

**Figure 2:**
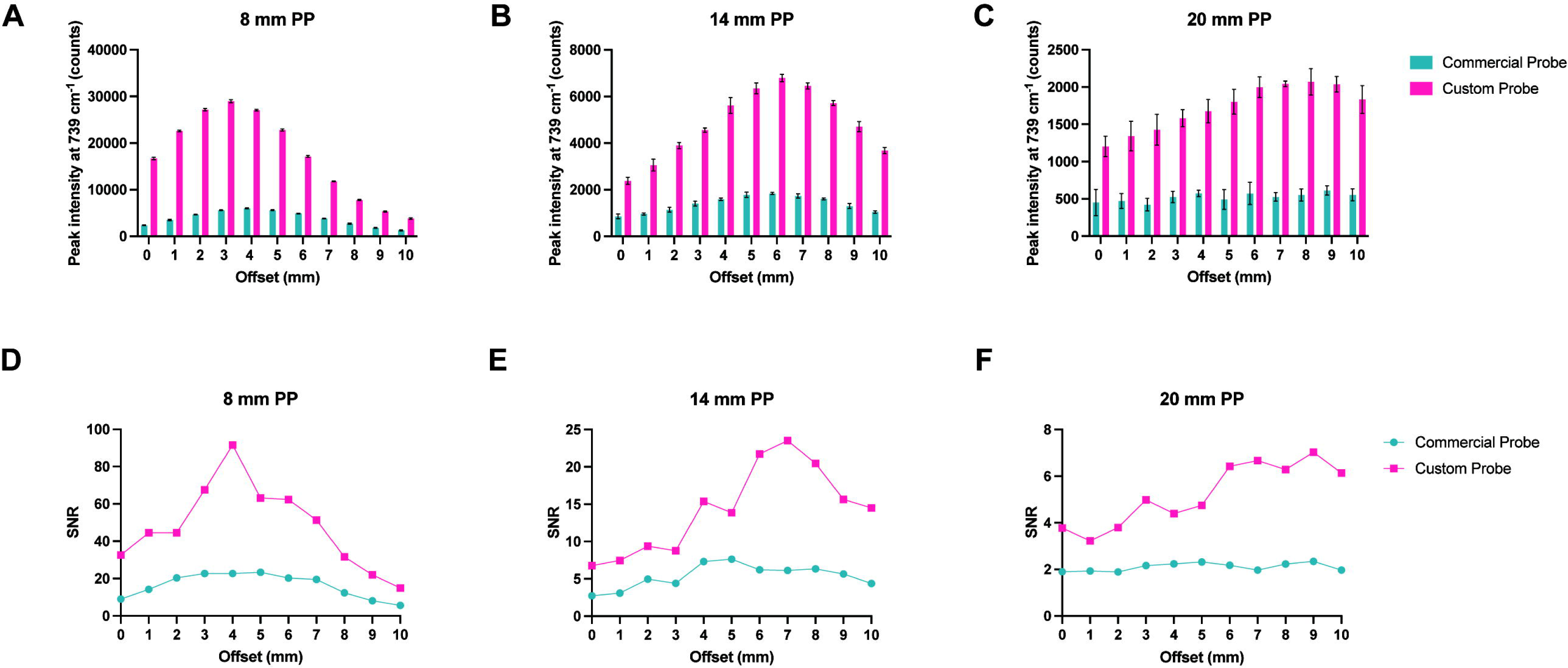
Comparison of the collection efficiencies of the commercially available collection probe and the custom-built collection probe. (A-C) Peak intensity of PTFE at 739 cm^−1^ through 8, 14, and 20 mm of polypropylene (PP) at 0 – 10 mm spatial offsets (1 mm increments). (D-F) Comparison of signal-to-noise ratios from each probe through 8, 14, and 20 mm of PP at 0 – 10 mm spatial offsets (1 mm increments). Spectra were acquired using a 4 s integration, 5 acquisitions, 785 nm laser, 500 mW laser power.

Having demonstrated the efficiency of our custom-built collection probe over the commercially available probe, we sought to reduce the overall time taken to sample an ROI by investigating the impact of sampling frequency on *in vivo* SESORRS imaging. Imaging approaches using conventional Raman approaches apply a tightly focused beam to achieve high resolution images, however, this technique typically requires time-intensive point-by-point acquisition of Raman spectra, hindering the real-time image acquisition desired for clinical applications. When imaging an ROI using Raman spectroscopy, the resulting image is not only a function of the laser spot size, but also of the distance between the sequentially scanned locations referred to as the sampling step or pixel size. There are three principal relationships between the laser spot and the sampling step: under-sampling, par-sampling, and over-sampling. Par-sampling, which involves matching the sampling step to the laser spot, provides an adequate estimation of the ROI, whereas over-sampling (where the step size is smaller than the laser spot size) has been shown to provide a significant improvement in image quality at the expense of total image acquisition time, given that the time taken to over-sample an ROI will be significantly higher than using a par-sampling approach on the same ROI [48–50]. In comparison to conventional Raman approaches, which typically utilize a laser beam with a spot size on the micron scale, SORS approaches typically utilize a laser beam with a spot size on the mm-cm scale [35]. Therefore, by taking advantage of sampling frequency approaches and increased laser-spot size and sampling frequency, we hypothesized that using a sampling frequency approach, it would be possible to image a significantly larger area in a shorter time frame in comparison to Raman approaches utilizing a tightly focused beam.

Sampling step size is determined by the laser spot size with a par-sampling frequency having the same steps size as the laser diameter in *x* and *y* and an over-sampling approach having a step size smaller than the laser diameter. Given that sampling frequency will influence the overall resolution of an image (i.e., a higher sampling frequency should delineate an ROI with greater certainty due to the acquisition of a greater number of data points from given area) we wanted to investigate how this could be applied to SESORRS imaging to reduce the overall time take to image an ROI. Calculations of the sampling frequency, or the number of spectral data points obtained per image, were performed based on the laser spot size of the elliptical beam, which measures 6 mm × 4 mm (**Figure 3A**). For par sampling, a step size of 6 mm in the x-direction and 4 mm in the y-direction is required. In the case of over-sampling by a factor of 10, the step sizes would need to be reduced to 0.6 mm in the x-direction and 0.4 mm in the y-direction. It is important to note that the sampling frequency directly impacts the overall resolution of the resulting image. In this study, we explored the effects of both par-sampling and over-sampling at frequencies of 2, 5, and 10, as illustrated in **Figure 3**, across a 12 mm × 12 mm area. Therefore, if we were to sample an area of 12 × 12 mm using each of the four sampling frequency approaches (par-sampling, over-sampling by 2, 5, and 10), we would need to perform a total of 12, 35, 176, and 651 measurements to create our image (**Figure 3B-E**). The step-size in *x* and *y* for each sampling frequency is summarized in **Table 1**. We hypothesized that in comparison to a par-sampling frequency approach, over-sampling by a small number (e.g., 2 and 5) would give rise to higher resolution images; however, further increases in the sampling frequency (e.g., 10) would not generate a significant increase in image contrast. Moreover, a higher sampling frequency would result in an increase of total time taken to image an ROI. Thus we sought to investigate the influence of sampling frequency approach on image quality.

**Figure 3:**
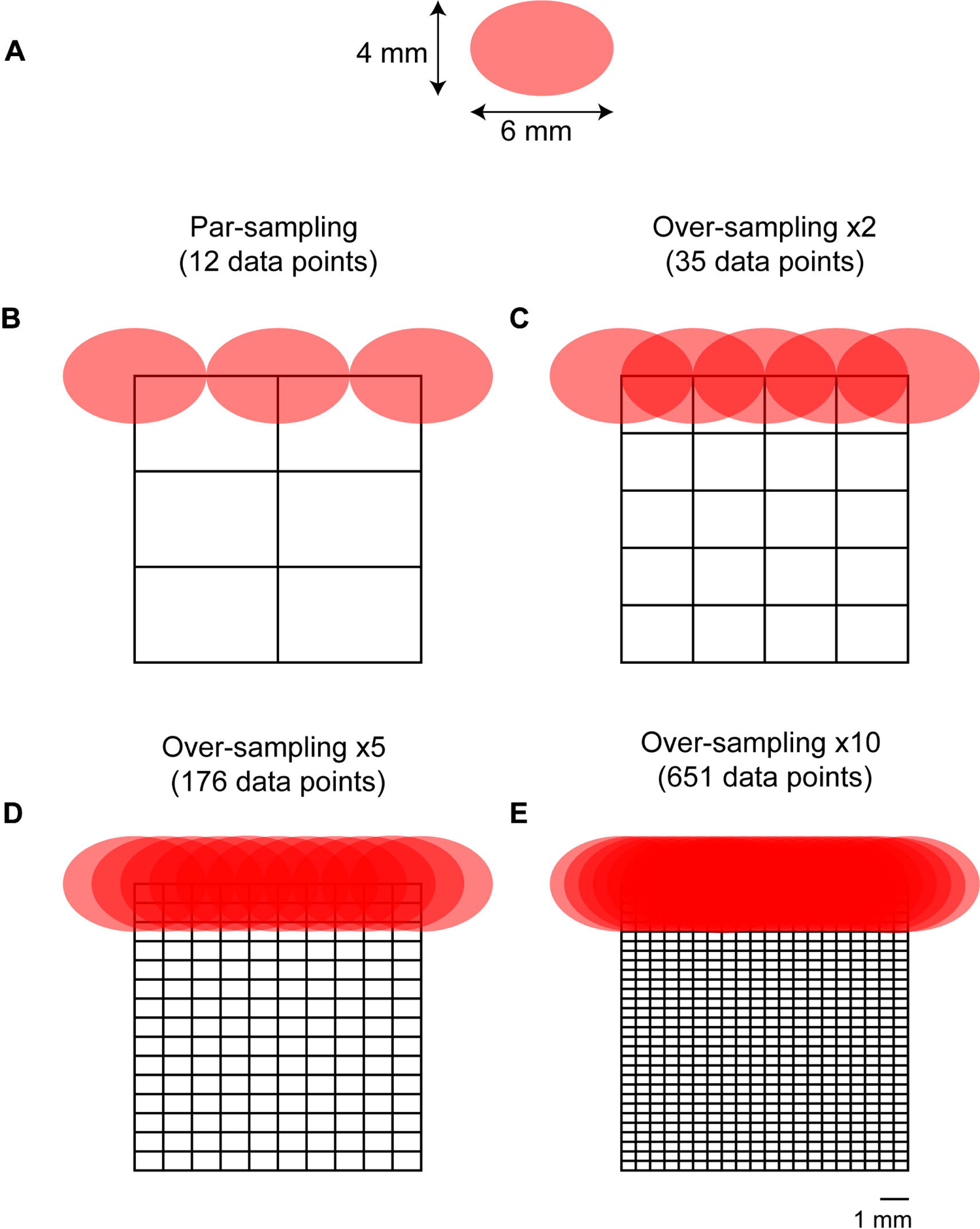
Diagram demonstrating a par-sampling and over-sampling by 2, 5 and 10 over a 12 × 12 mm area. (A) laser spot-size. The beam is elliptical and thus has a diameter of 6 mm × 4 mm. (B-E) diagram demonstrating the number of measurements per sampling frequency of a 12 × 12 mm area. Par sampling represents a step size of 6mm in × and 4mm in y, whereas a x10 sampling frequency represents a 0.6mm step size in *x* and a 0.4mm step size in *y*. Stepwise measurements are performed in *x* and then in *y* to image the whole 12 × 12 mm area. (B) Par-sampling approach resulting in a total of 12 data points collected over the ROI. (C) Over-sampling by 2 resulting in a total of 35 data points collected over the ROI. (D) Over-sampling by 5 resulting in a total of 176 data points collected over the ROI. (E) Over-sampling by 10 resulting in a total of 651 data points collected over the ROI.

To investigate the influence of sampling frequency on image quality, we prepared a brain-tumor phantom that was representative of an *in vivo* GBM. Briefly, the brain of a 12 week old-euthanized mouse was removed, and the skin over the face of the mouse left intact. The head was then fixed in 4% PFA and then transferred to 70% EtOH. PTFE (5 mm × 5 mm, thickness 2 mm) was glued directly underneath the skull to create a brain tumor mimic (**Figure 4A**) and then filled with agarose gel. Unlike previous work where the skin of the mouse was removed from the head and we wrapped the skull in porcine tissue to create a skin mimic, only fur was removed using hair removal cream and the skin was left on the mouse’s head (**Figure 4B**). The mouse head was placed on a paraffin wax mold (**Figure 4B**) to allow repeated measurements of the head over multiple days by ensuring the mouse head was continually placed in the same position as the prior day. The mapping of PTFE through the mouse head was then carried out at spatial offset of 1-3 mm, 0.5 mm increments. The spatial offset was controlled by moving the *xyz* translational stage away from the point of collection. The spatial offset kept fixed for each set of measurements, and the sample was moved using the *xy* stage (i.e., the excitation and collection optics remained static at each spatial offset). Measurements were taken at varying step sizes depending on the sampling frequency being used (**Table 1**). Surface/contour false color 2D heat maps were then constructed using the peak height of PTFE (739 cm^−1^).

**Figure 4:**
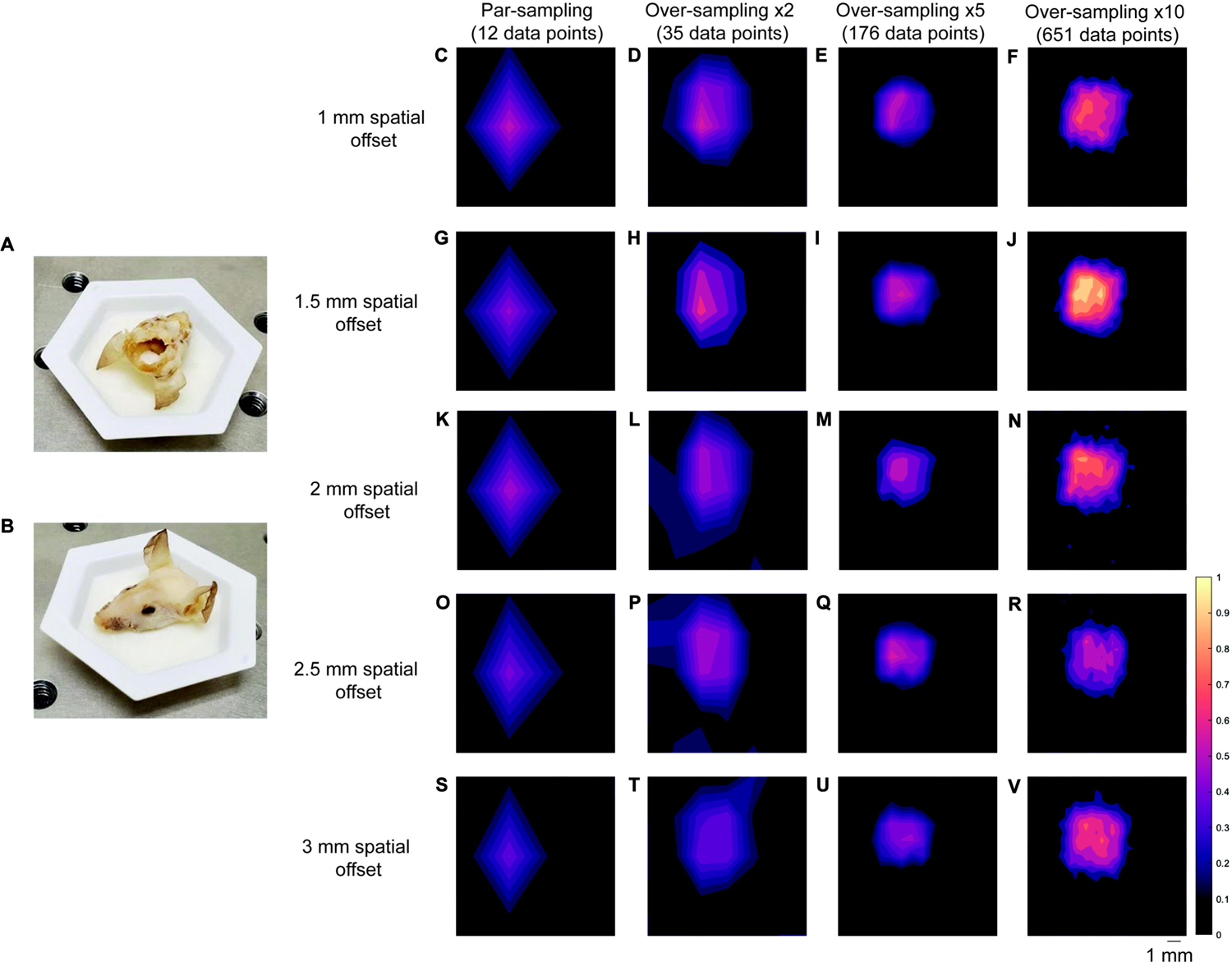
Application of varying sampling frequency approaches (par, x2, x5 and x10) in brain tumor phantoms at 1-3 mm spatial offsets (0.5 mm increments). (A) PTFE placed under the mouse’s skull. (B) Mouse head placed in paraffin wax mold. (Figure 4C-F) Mapping of the brain tumor phantom at using a par-sampling frequency and oversampling by 2, 5, and 10 at a 1 mm spatial offset. (G-J) Mapping of the brain tumor phantom using a par-sampling frequency and oversampling by 2, 5, and 10 at a 1.5 mm spatial offset. (K-N) Mapping of the brain tumor phantom using a par-sampling frequency and oversampling by 2, 5, and 10 at a 2 mm spatial offset. (O-R) Mapping of the brain tumor phantom using a par-sampling frequency and oversampling by 2, 5, and 10 at a 2.5 mm spatial offset. (S-V) Mapping of the brain tumor phantom using a par-sampling frequency and oversampling by 2, 5, and 10 at a 3 mm spatial offset. All measurements were acquired using the custom-built collection probe, 785 nm laser, 0.5s integration time, 1 accumulation.

**Figure 4** demonstrates the mapping of the brain tumor phantom using each of the four sampling frequencies at spatial offsets of 1mm (**Figure 4C-F**), 1.5 mm (**Figure 4G-J**), 2 mm (**Figure 4K-N**), 2.5 mm (**Figure 4O-R**), and 3 mm (**Figure 4S-V**). In this case, over-sampling by 10 at a 1.5 mm spatial offset gave rise to the highest signal, thus all other images were normalized to the most intense peak in this image. As shown in **Figure 4**, when the sampling frequency is increased, the ability to delineate the PTFE square is also improved. As anticipated, the difference in image quality between images acquired using a sampling frequency of 5 and 10 is subtle, indicating that there may be limited advantages to using a higher sampling frequency approach. Encouragingly, the area of intensity that corresponds to the PTFE signal is in agreement with the size of PTFE that was glued beneath the mouse’s skull (5 mm × 5 mm). In previous work, we demonstrated the detection of PTFE through a brain tumor where the skin was removed from the mouse’s head, PTFE embedded underneath the skull and the head then wrapped in 2 mm of porcine tissue and imaged using a SORS approach at 2, 2.5, and 3 mm spatial offset, 1 mm step sizes [33]. In comparison to our previous work, by over-sampling by 5 and by 10 we demonstrate improved delineation of PTFE at 1, 1.5, 2-, 2.5-, and 3-mm spatial offsets. In this work, even at lesser sampling frequencies (i.e., par and 2), we demonstrate successful detection of a rough ROI and therefore hypothesize that we could use such sampling frequency to scan large areas of tissue, identity ROIs, and then delineate these areas with greater certainty using a larger sampling frequency approach *in vivo*. Our data indicate that a spatial offset of 1.5 mm is particularly suitable for *in vivo* SESO(R)RS imaging of GBM using our custom-built collection optics. This offset effectively suppresses surface signals, as evidenced by the reduction in spectral contributions from tissue and bone peaks at 1440 cm^−1^ and 957 cm^−1^, respectively. Moreover, it produces a strong contribution from PTFE at 739 cm^−1^, resulting in both high SNRs and intense signal. In our decision to proceed with a 1.5 mm spatial offset, we are confident that any of the five spatial offsets under consideration would have been apt for *in vivo* imaging. Furthermore, considering that the selection of spatial offset is contingent on the specific sample, and that the depth of an analyte frequently remains unknown, our findings indicate that it is feasible to capture images of a ROI at different spatial offsets without substantially affecting the overall image quality. This capability is especially valuable for *in vivo* imaging, where tumor growth is inherently three-dimensional and the extent of tumor invasion may not be comprehensively characterized before imaging.

Building on our proof-of-concept measurements using brain tumor phantoms, we then proceeded with *in vivo* studies using SERRS CAs (**Figure 5A**). SERRS CAs were synthesized using gold nanostars which were functionalized with a resonant Raman reporter molecule (IR780p) and encapsulated in a silica shell. The silica shell was then functionalized with PEG_5000_ to increase their biocompatibility (**Figure 5A**). For this work, we chose a commercially available near-IR active dye (IR780p) due to its resonance properties. Such properties ensure the greatest enhancement of Raman signal following interaction of the SERRS CAs with incident light (785 nm). The corresponding SERRS spectrum of the SERRS CAs is shown in **Figure 5B**. SERRS CAs were also characterized by zeta potential analysis, nanoparticle tracking, and TEM (**Figure 5C**). The final SERRS Cas used in this study had a diameter of 133 ± 12 nm, as confirmed by TEM. The zeta potential of the SERRS CAS was −48 mV after thiolation and −15 mV after pegylation indicating successful functionalization of SERRS CAs with PEG_5000_. Following synthesis, SERRS CAs were injected via the tail vein to GBM tumor-bearing BALB/cJ mice. Tumor incidence and size was confirmed by MRI at 2 weeks post intracranial injection of GL261-Lu cells, **Figure 6A-C**. In this instance SERRS CAs were not functionalized with a targeting ligand, thus we relied solely on the EPR effect to ensure uptake of the NPs at the ROI. Previous work from our group has demonstrated the successful targeting of various solid tumors *in vivo* using targeted SERRS CAs; thus, the purpose of this work was not to evaluate potential targeting ligands, but to demonstrate the potential of SESORRS imaging for the detection and delineation of ROIs *in vivo* and improve the overall collection efficiency of our approach. In our previous report using SESORRS for the imaging of GBM, we used a SERRS CA concentration of 8 nM [33]. In this work, however, due to optimization of the NP synthesis protocol from previous reports and optimization of the collection probe, we were able to reduce the concentration of the injected dose to 2 nM, demonstrating a significant improvement on previous reports on the use of SERRS CAs for *in vivo* imaging using SESORRS [33]. Biodistribution studies on the uptake of SERRS CAs *in vivo* confirmed the accumulation of NPs in the liver, spleen, and lymph nodes (**Figure S3**) which is consistent with other reports involving SERRS nanoparticles [15,16,29,33,51].

**Figure 5:**
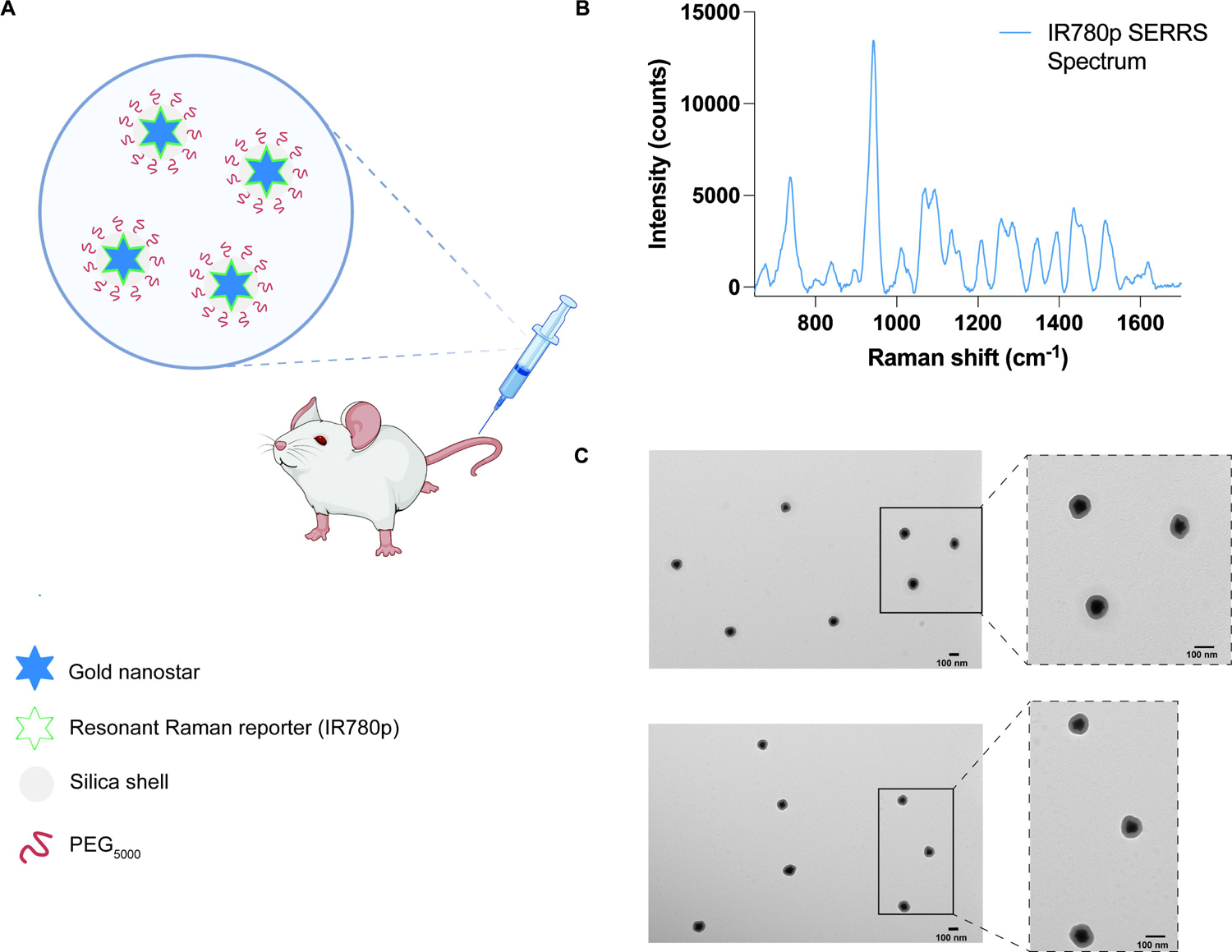
Characterization of SERRS nanostars for *in vivo* administration. (A) Conceptual figure showing gold nanostars administered via the tail vein. Gold nanostars were functionalized with a resonant Raman reporter molecule (IR780p) and then encased in a silica shell which was then functionalized with PEG_5000_. (B) The unique fingerprint spectrum of the SERRS nanostars functionalized with IR780p. The spectrum corresponds to that of IR780p. SERRS spectra were obtained using a 785 nm wavelength, 100 ms integration time. (C) Transmission electron microscope image of the PEGylated SERRS nanostars from two different areas on the TEM grid. The scale bar represents 100 nm. Together the nanostars and silica shell had a total average diameter of 133 ± 12 nm. Images were acquired using a JEOL 1200EX Transmission Electron Microscope at 80 kV.

**Figure 6:**
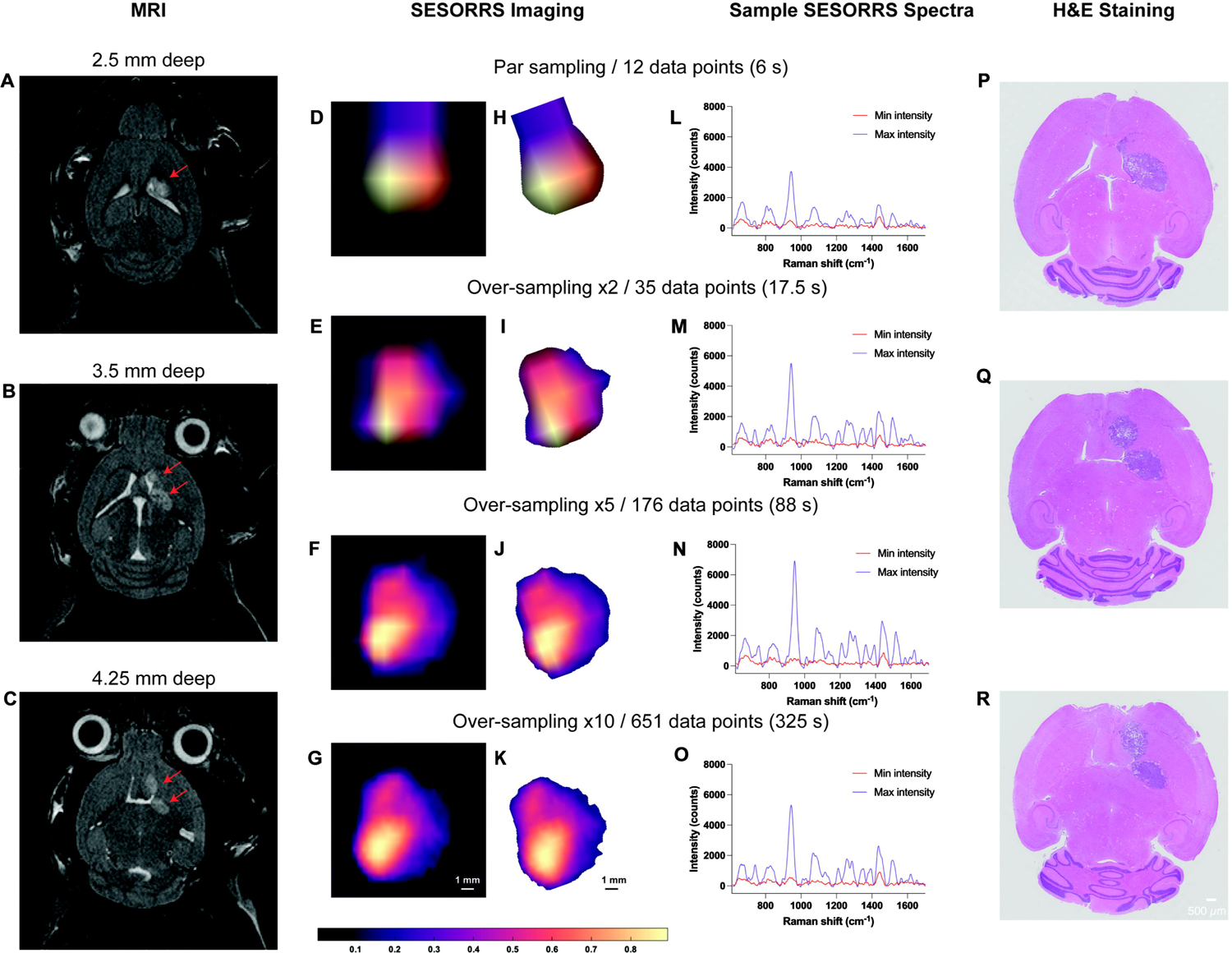
Detection of GBM in mice (n=3) using SESORS and optimized sampling approaches. (A-C) Magnetic resonance imaging at 2 weeks post-injection of GL261-Luc cells confirms the presence of a tumor as indicated by the red arrows. Magnetic resonance images acquired at depths of 2.5 mm, 3.5 mm, and 4.25 mm respectively. MR images demonstrate the emergence of a secondary tumor region at greater depths within the brain. Following confirmation of tumor growth, non-targeted SERRS NPs were administered to tumor-bearing mice (2nM, 100µL) via the tail vein. *In vivo* SESORRS imaging was performed using a spatial offset of 1.5mm, using (D) par sampling frequency (12 data points) (E) over-sampling frequency of: 2 (35 data points), (F) 5 (176 data points), and (G) 10 (651 data points). The respective times taken to acquire each image are also shown. Raman spectra were truncated, normalized, and principal component (PC) analysis was applied to generate false-color 2D heat maps. PC scores 2 and 3 were used to create SESORS images and represent the accumulation of SERRS NPs in the tumor. SESORRS measurements were acquired using a power density of 6.5 mW/mm2, 1.5mm spatial offset, 0.5s integration time, 1 acquisition, 785 nm excitation wavelength. (H-K) Background removal of areas of non-intensity from the SESORRS images. Images were then rotated to match the orientation of the MR images. (L-O) SESORRS spectra taken at the point of minimum and maximum intensity from the respective SESORRS image using a par sampling frequency and an oversampling frequency of 2, 5 and 10 respectively. (P-R) H&E stained 5µM section of the brain confirming the presence of a tumor and the emergence of a secondary tumor region as deeper regions are sliced.

The application of varying sampling frequency approaches to the imaging of GBM through the intact skull using a SESORRS approach is described in **Figure 6**. MRI confirms the presence of a left frontal tumor at a depth of 2.5 mm and, interestingly, the emergence of a secondary ROI at deeper depths is also observed via MRI at depths of 3.5 and 4.25 mm (**Figure 6B** and **C** respectively). Subsequent SESORRS imaging was then performed using a par-sampling frequency and over-sampling frequencies of 2, 5, and 10 over the same area on each mouse (**Figure 6D-G**). Images were constructed following application of PCA and principal components 2 and 3 were folded to a spectral image. The total acquisition time (6 s, 17.5 s, 88 s, and 325 s) at each frequency is also indicated. Background removal of areas of non-intensity were then subtracted from the SESORRS images (**Figure 6H-K**) and the images were then rotated anti-clockwise (−20°) to match the orientation of the magnetic resonance (MR) images. As sampling frequency increases the ability to delineate the ROI is increased, however there is little difference in resolution between the images acquired using an over-sampling frequency of 5 and 10, thus suggesting that such extreme sampling frequencies are not necessary for in vivo SESORRS imaging. In addition to improved image resolution, the emergence of a secondary ROI with lower spectral intensity is also observed at all four sampling frequencies, indicating that our SESORRS approach is able to detect deeper seated lesions and also to detect signal that is proportional to depth. This is in agreement with the theory of photon migration in tissue (i.e., deeper born photons will give rise to lower intensity signal). Sample spectra taken at points of maximum and minimum intensity from the respective image using a par-sampling frequency and an over-sampling frequency of 2, 5 and 10 respectively are also shown (**Figure 6L-O**). In all four cases, the data clearly demonstrate the detection of SERRS CAs in the pixels of maximum intensity due to detection of several peaks related to IR780p including the main peak at 947 cm-1 which is attributed to IR780p (i.e., the resonant Raman reporter used in this study). Spectra taken at the point of minimum intensity show spectral contribution solely from tissue (e.g., 1440 cm-1), indicating selective uptake of the SERRS CAs within the tumor and not in healthy tissues. *Ex vivo* H&E staining also confirmed the presence of a tumor and emergence of a secondary tumor region as deeper regions of the brain are sliced (**Figure 6P-R**). Moreover, the histopathological findings were also in agreement with the MR images (**Figure 6A-C**).

The main aim of this study was to develop a more efficient SORS imaging system and approach in order to reduce the overall time take to image a ROI. In our previous report, we used a 15s integration time, (3s, 5 acquisitions) which resulted in lengthy image acquisition at each data point. In this work however, we utilized a 0.5s integration time and 1 acquisition at each data point, demonstrating a significant improvement in the collection efficiency of our custom-built SORS collection probe (30-times shorter integration time per acquisition point). The ability to map the whole head of the mouse and determine a specific region of interest using as few as 12 spectra (total acquisition time 6s) is achieved and we demonstrate that the subsequent use of a higher sampling frequency can delineate tumor margins in the ROI with greater certainty (**Figure 6D-K**). Similarly, in comparison to our previous report, we have also been able to lower the integration time used for the detection of GBM using a SESORRS approach from 13.8 mW/mm2 to 6 mW/mm2 [33]. We believe this to be within the MPE limit however we point out that MPEs are set for accidental exposure to radiation and not for intentional exposure, e.g., for use in a clinical procedure. Therefore, the clinical limit could, and most likely would, differ when considering the benefit to the patient.

In addition to lowering the overall time taken to image an ROI, and perhaps most importantly, this work demonstrates the possibility to delineate a secondary ROI deeper within the brain using our SESORRS approach at all four sampling frequencies. In this case, the development of tumor grown in the deeper-seated area was spontaneous, meaning we did not purposely create two regions through the intracranial injection of mice with GL261-Luc cells. This is representative of an *in vivo* system in which tumor growth is spontaneous and heterogenous in nature. With this in mind, we believe that when looking to the future, SESORRS imaging could be applied for the intra-operative assessment of ROIs using this sampling frequency approach and may offer a benefit over current image guided surgical approaches. Following tumor resection, it is well established that microscopic malignant lesions are often missed by visual inspection. Currently, fluorescence probes lead the way in image guided surgical applications [4], however, both fluorescence and traditional SERRS approaches are only able to image superficial lesions of interest. Moreover, fluorescence imaging suffers from photobleaching and intrinsic autofluorescence [8]. Surgical resection is stopped once the surgeon is confident all malignant lesions have been removed, however there is a possibility that deep-seated, microscopic lesions remain due to the inability of fluorescence imaging to detect deep-seated lesions. Thus, we believe that it could be possible to apply SESORRS imaging using a par-sampling frequency to scan large areas of tissue to identify ROIs where the malignant tissues are present. Then, using a higher-sampling frequency approach, e.g. 2 or 5, delineate that same ROI with greater certainty and identify the presence of potentially deeper seated lesions in real time. Future work will therefore focus on developing this approach through the incorporation of a white light camera to correlate what is seen by the naked eye with the corresponding SESORRS image, evaluating the efficiency of SESORRS–image guided surgery for the detection of deeper-seated lesions, i.e., determining signal contribution relative to depth *in vivo* using a multi-modal imaging approach. In this instance a spatial offset of 1.5 mm was used for SORS imaging, however it is reasonable to assume that the other spatial offsets would have been suitable for SESORS imaging, e.g., 1, 2 and 2.5 mm. This is of course dependent on the sample under study it is reasonable to assume that the use of larger spatial offsets may lead to the detection of even deeper-seated ROI however this would come at the expense of spectral resolution and may require longer integration times.

## CONCLUSION

Through improvements in the collection efficiency of our imaging probe, and consideration of sampling frequency approach, we demonstrate the ability to detect image GBM through the intact skull using a SESORRS approach. We demonstrate that in comparison to previous reports, the ability to identify and delineate a ROI can in fact be carried out relatively quickly and does not rely on such time-intensive measurement. To the best of our knowledge, no one has applied this approach for the *in vivo* imaging of cancer using conventional Raman, SERRS, SORS or SESORRS approaches. Importantly our results demonstrate the ability to detect deeper-seated tumors in agreement with MRI and histopathology in a much shorter time-frame than what has been previously reported. Moreover, the work presented here is the first to identify a secondary ROI using a SESORRS approach for the detection of cancer *in vivo*. Unlike other optical imaging approaches, SESORRS offers the opportunity to image deeper into tissue. We believe that the approach outlined here opens the potential for SESORRS to be applied in image-guided applications for the detection of multiple solid tumor types and demonstrates an important step forward in the development and application of SESORRS imaging for the detection of cancer *in vivo*.

## ACKNOWLEDGEMENTS

This work was supported by the following grants: DFCI Trustee Science Committee Postdoctoral Fellowship, Claudia Adams Barr Award for Innovative Cancer Research (DFCI), and K99CA266921 to F.N.; an award from the Cancer Research UK Grand Challenge and the Mark Foundation to K.M.H. and the SPECIFICANCER team; TAMU and TEES start-up funding, NSF-Engineering Research Center for Precise Advanced Technologies and Health Systems for Underserved Populations (PATHS-UP)-Award Number 1648451, and NSF Funding Award Number: 2022805 to S.M. We also thank the pre-clinical animal imaging core facility (Lurie Family Imaging Center, Dana-Farber Cancer Institute) for technical assistance and advice, and acknowledge the initial contributions of Moritz F. Kircher, MD, PhD (now deceased) to the work presented here. Figures 1 and 5 were created in part, using Biorender.com.

## CONFLICTS OF INTEREST

SR has several filed patents in the areas of Wavelength Stabilized Lasers, Raman Probes, Raman Concatenation, dual wavelength lasers for fluorescence mitigation and fluid analysis using Raman spectroscopy.

## Notes

### Competing Interest Statement

Scott Rudder has several filed patents in the areas of Wavelength Stabilized Lasers, Raman Probes, Raman Concatenation, dual wavelength lasers for fluorescence mitigation and fluid analysis using Raman spectroscopy.

