## Supplementary information for "In vivo Imaging using Surface Enhanced Spatially Offset Raman Spectroscopy (SESORS): Balancing Sampling Frequency to Improve Overall Image Acquisition"

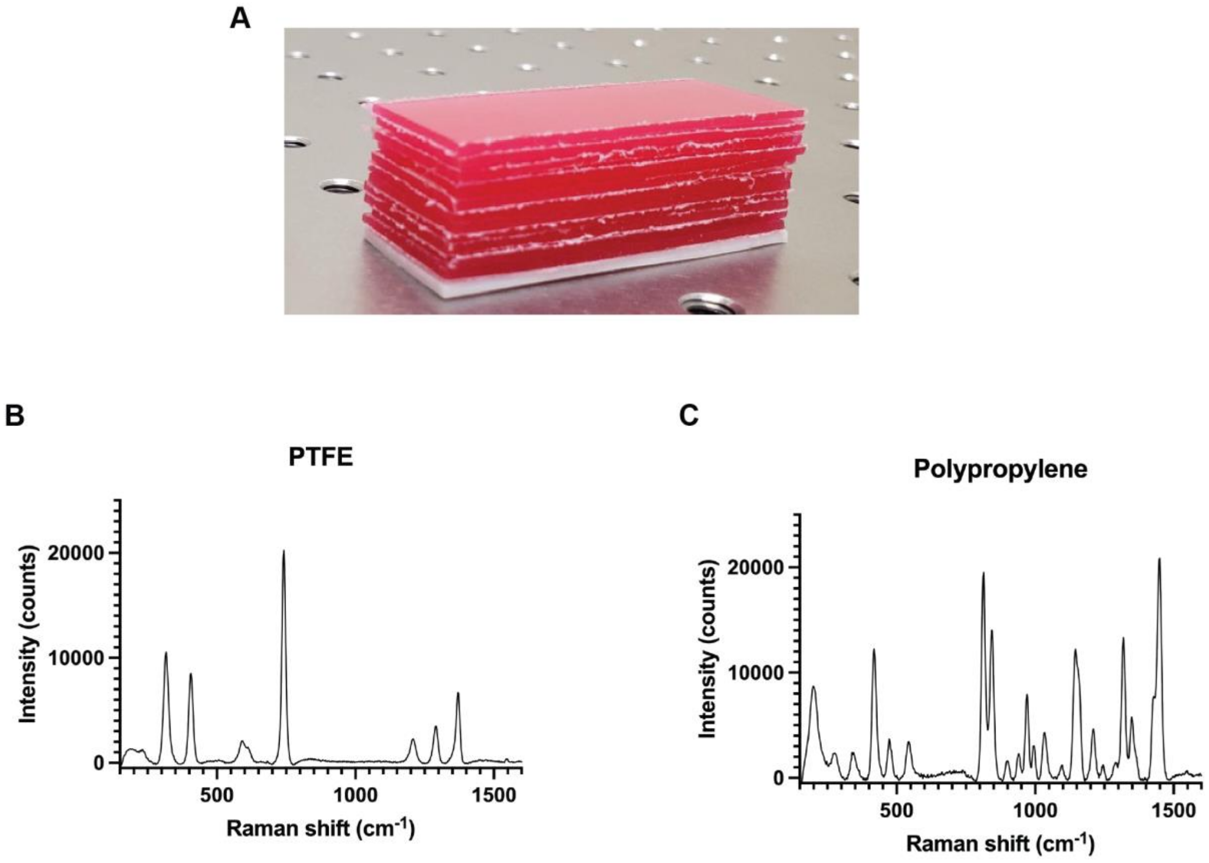


**Figure S1: Calibration standards and their corresponding Raman spectrum. (A) Sheets of pink polypropylene (PP) were placed on top of white polytetrafluoroethylene (PTFE) to create calibration standards. (B) Raman spectrum of PTFE. (C) Raman spectra of PP. Spectra were acquired using a 785 nm laser, 1s integration time.**

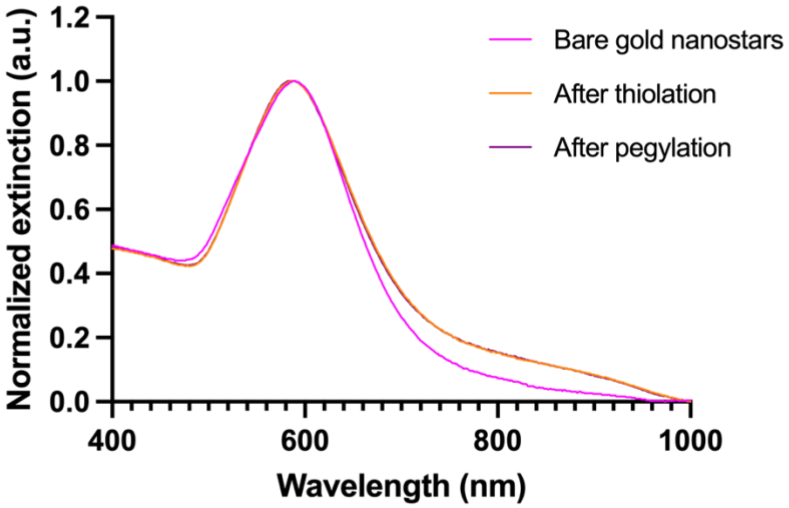


**Figure S2: Extinction spectroscopy of SERRS CAs after thiolation and after PEGylation. Spectra were characterized using a UV-2600 UV-VIS spectrophotometer (Shimadzu).**

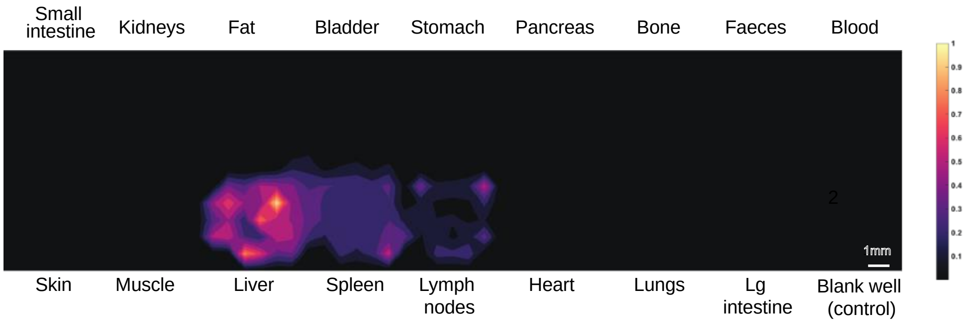


**Figure S3: Biodistribution of SERRS CAs in tissues. GL261 tumor-bearing mice (n=3) were injected with 100 μL of 2 nM of SERRS nanostars, and tissues of interest harvested and homogenized. The tissues were then analyzed by Raman imaging (25% laser power, 785 nm laser, 1 s acquisition time, 5× objective) to determine the relative accumulation in different tissues. Images are representative of n = 3 mice, 1 organ per well. Measurements were acquired using a LabRAM HR Evolution (HORIBA Scientific).**
